# Crucial role for Sodium Hydrogen Exchangers in SGLT2 inhibitor-induced arterial relaxations

**DOI:** 10.1101/2023.12.05.570303

**Authors:** Elizabeth A Forrester, Miguel Benítez-Angeles, Kaitlyn E. Redford, Tamara Rosenbaum, Geoffrey W Abbott, Vincenzo Barrese, Kim Dora, Anthony P Albert, Johs Dannesboe, Isabelle Salles-Crawley, Thomas A Jepps, Iain A Greenwood

## Abstract

1

**Introduction:** Sodium dependent glucose transporter 2 (SGLT2 or SLC5A2) inhibitors effectively lower blood glucose and are also approved treatments for heart failure independent of raised glucose. One component of the cardioprotective effect is reduced cardiac afterload but the mechanisms underlying peripheral relaxation are ill defined and variable. We speculated that SGLT2 inhibitors promoted arterial relaxation via the release of the potent vasodilator calcitonin gene-related peptide (CGRP) from sensory nerves independent of glucose transport.

**Experimental approach:** The functional effects of SGLT2 inhibitors (dapagliflozin, empagliflozin, ertugliflozin) and the sodium/hydrogen exchanger 1 (NHE1) blocker cariporide were determined on pre-contracted mesenteric and renal arteries from male Wistar rats using Wire-Myography. SGLT2, NHE1, CGRP and TRPV1 expression in both arteries was determined by Western blot and immunohistochemistry. Kv7.4/5/KCNE4 and TRPV1 currents were measured in the presence and absence of dapagliflozin and empagliflozin.

**Results:** All SGLT2 inhibitors produced a concentration dependent relaxation (1µM-100µM) of mesenteric arteries that was considerably greater than in renal arteries. Cariporide relaxed mesenteric arteries but not renal arteries. Immunohistochemistry with TRPV1 and CGRP antibodies revealed a dense innervation of sensory nerves in mesenteric arteries that was absent in renal arteries. Consistent with a greater sensory nerve component, the TRPV1 agonist capsaicin produced significantly greater relaxations in mesenteric arteries compared to renal arteries. Relaxations to dapagliflozin, empagliflozin and cariporide were attenuated by incubation with the CGRP receptor antagonist BIBN-4096, the Kv7 blocker linopirdine and the TRPV1 antagonist AMG-517 as well as by depletion of neuronal CGRP. Neither dapagliflozin nor empagliflozin directly activated heterologously expressed TRPV1 channels or Kv7 channels. Strikingly, only NHE1 colocalised with TRPV1 in sensory nerves, and cariporide pre-application prevented the relaxant response to SGLT2 inhibitors.

**Conclusions:** SGLT2 inhibitors relax mesenteric arteries by a novel mechanism involving the release of CGRP from sensory nerves following inhibition of the Na^+^/H^+^ exchanger.

## 2 Introduction

Inhibitors of sodium-dependent glucose transporter 2 (SGLT2 encoded by *SLC5A2*), such as dapagliflozin, empagliflozin or canagliflozin, lower blood glucose levels through increased urinary excretion of glucose (John et al. 2016). These drugs are also effective treatments for heart failure independent of raised glucose with the UK National Institute for Health and Care Excellence (NICE) recommending them for treatment of heart failure with reduced ejection fraction (McMurray et al. 2019; Nassif et al. 2019; Wheeler et al. 2021). Various mechanisms have been suggested for the beneficial cardiac effects of SGLT2 inhibitors including decreased cardiac fluid retention, reduced reactive oxygen species generation, and lessened fibrosis allied to a decrease in afterload through arterial relaxation. The ability of SGLT2 inhibitors to suppress cardiac fibrogenesis and oxidative stress mimics the effect of blockers of the sodium/hydrogen exchanger 1 (NHE1) such as cariporide (Baartscheer et al. 2017; Zhang et al. 2021; Uthman et al. 2022; Chung et al. 2023) and studies have shown that SGLT2 inhibitors also block NHE isoforms (Baartscheer et al. 2017; Uthman et al. 2018; De Stefano et al. 2021) with in silico studies predicting a binding site in the extracellular sodium-binding pocket of NHE1 (Uthman et al. 2018; Iborra-Egea et al. 2019). In contrast to its effects in the heart, relatively little is known about the effects of SGLT2 inhibitors in the vasculature. SGLT2 inhibitors do lower blood pressure and directly relax aorta and mesenteric arteries ex vivo (Li et al. 2018; Seo et al. 2020; Hasan and Hasan 2021; Hasan, Menon, et al. 2022; Hasan, Zerin, et al. 2022) but the underlying mechanisms are ill defined with variable contributions from voltage-dependent potassium (Kv) channels, endothelium and protein kinase G implicated. Resistance arteries that dictate blood pressure are richly innervated with peptidergic sensory nerves (Aalkjaer et al. 2021). Release of calcitonin-gene related peptide (CGRP) from sensory nerves has a pronounced vasodilatory effect in many arteries (Brain et al. 1985; Shiraki et al. 2000, 2001). Sensory nerve terminals contain several ion channels (including Transient Receptor Potential ion channels TRPV1, TRPA1 and TRPM3) that respond to lipid mediators, low pH, and noxious chemicals (e.g., capsaicin, allicin) to cause CGRP release (Aalkjaer et al. 2021). Kawasaki et al (2009) suggested that protons released contemporaneously with noradrenaline from sympathetic nerves can activate TRPV1 in mesenteric arteries. There is also evidence for sensory nerve-endothelium cross talk (Shiraki et al. 2001). We speculated that the vasodilatory effect of SGLT2 inhibitors was manifest by an action on sensory nerves and the concomitant release of CGRP, underpinned by NHE1 modulation.

## 3 Methods

### 3.1 Animals

Experiments were performed on arteries from male rats aged 11-14 weeks and weighing 175-300g, housed within the Institute of Cancer Research facility (Surrey, UK). Rats were housed in cages with LSB Aspen woodchip bedding and a 12-hour light/dark cycle, maintained at constant temperature (21±1°C) and humidity (50%±10%), with free access to water and food, all in accordance with the Animal Scientific Procedures Act 1986 (ASPA). The tissues were transported each day to St George’s University of London, bathed and preserved in a salt solution containing (mmol·L^-1^): 126 NaCl, 6 KCl, 10 Glucose, 11 HEPES, 1.2 MgCl_2_ and set to pH7.4 with NaOH. Animals were sacrificed by cervical dislocation and death confirmed via severing the femoral artery circulation in accordance with Schedule 1 of the ASPA 1986.

Second order mesenteric and the left and right main renal arteries were dissected, cleaned of fat and adjacent tissue and stored on ice in physiological salt solution (PSS) containing (mmol·L^-1^): 119 NaCl, 4.5 KCl, 1.17 MgSO_4_·7H_2_0, 1.18 NaH_2_PO_4_, 25 NaHCO_3_, 5 glucose, 1.25 CaCl_2_.

### 3.2 Cell Cultures

HEK293 cells (ATCC® CRL-1573) were cultured in a complete growth medium containing Dulbecco’s modified Eagle’s medium with high-Glucose (DMEM GibcoTM) complemented with 10% fetal bovine serum (HyCloneTM) and 100 U/ml of Penicillin-Streptomycin (GibcoTM). Cell cultures were maintained in a humidified incubator at 37°C with an atmosphere of 95% of air and 5% of CO_2_. Cells were subcultured every 3 days using 0.25% (w/V) Trypsin-EDTA solution (GibcoTM).

### 3.3 Electrophysiology

The effect of various SGLT2 inhibitors was assessed on currents generated by the over-expression of Kv7 and TRPV1 genes in *Xenopus laevis* oocytes and HEK293 cells respectively.

#### 2.3.1 Channel subunit cRNA preparation and *Xenopus laevis* oocyte injection

We generated cRNA transcripts encoding human Kv7.4, Kv7.5 and KCNE4 by in vitro transcription using the mMessage mMachine kit (Thermo Fisher Scientific, Waltham, MA, USA) according to manufacturer’s instructions, after vector linearization, from cDNA sub-cloned into expression vectors (pTLNx and pXOOM) incorporating *Xenopus laevis* β-globin 5’ and 3’ UTRs flanking the coding region to enhance translation and cRNA stability. We injected defolliculated stage V and VI *Xenopus laevis* oocytes (Xenoocyte, Dexter, MI, USA) with Kv7 and KCNE4 cRNAs (2-10 ng) and incubated the oocytes at 16 °C in ND96 oocyte storage solution containing penicillin and streptomycin, with daily washing, for 2-5 days prior to two-electrode voltage-clamp (TEVC) recording.

### 3.4 Two-electrode voltage clamp (TEVC)

We performed TEVC at room temperature using an OC-725C amplifier (Warner Instruments, Hamden, CT, USA) and pClamp10 software (Molecular Devices, Sunnyvale, CA, USA) 2-5 days after cRNA injection. We placed oocytes in a small-volume oocyte bath (Warner) and viewed them with a dissection microscope for cellular electrophysiology. We sourced chemicals from Sigma-Aldrich (St. Louis, MO, USA). We studied the effects of SGLT2 inhibitors, solubilized in bath solution (in mM): 96 NaCl, 4 KCl, 1 MgCl2, 1 CaCl2, 10 HEPES (pH 7.6). Compounds were introduced into the oocyte recording bath by gravity perfusion at a constant flow of 1 ml per minute for 3 minutes prior to recording. Pipettes were of 1-2 MΩ resistance when filled with 3 M KCl. Currents were recorded in response to voltage pulses between −80 mV and +40 mV at 20 mV intervals from a holding potential of −80 mV, to yield current-voltage relationships. Data were analyzed using Clampfit (Molecular Devices) and Graphpad Prism software (GraphPad, San Diego, CA, USA), stating values as mean ± SEM. Raw tail currents were plotted versus prepulse voltage and fitted with a single Boltzmann function:

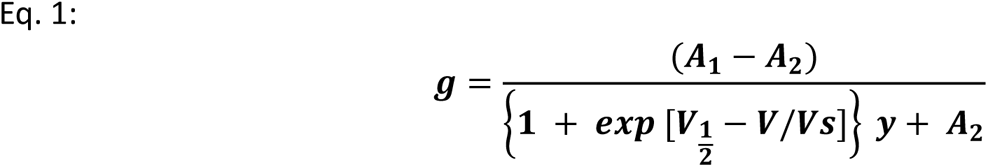

where g is the normalized tail conductance, A_1_ is the initial value at -∞, A_2_ is the final value at +∞, V_1/2_ is the half-maximal voltage of activation and Vs the slope factor.

#### 2.4.1 Transient cell transfection and patch-clamp

For patch-clamp experiments, the cells were seeded on poly-D-lysine (1 mg/ml, Sigma) coated coverslips for patch-clamp experiments. After 24 hrs, HEK293 cells were transfected with plasmids pcDNA3.1 containing the rat wild-type TRPV1 channel (1.5 µg) and pcDNA3.1 with green fluorescent protein (GFP, 500 ng) to identify successfully transfected cells using jetPEI™ Polyplus transfection reagent, per manufacturer’s instructions.

TRPV1 currents were recorded from transiently transfected HEK-293 using the patch-clamp technique in the outside-out configuration (Hamill et al., 1981). Solutions were changed with an RSC-200 rapid solution changer (Molecular Kinetics). Currents were low-pass filtered at 2 kHz and sampled at 10 kHz with an EPC 10 amplifier (HEKA Elektronik) and were plotted and analysed with Igor Pro (Wavemetrics Inc.). Intracellular and extracellular recording solutions contained (in mM): 130 NaCl, 3 HEPES (pH 7.2) and 1 EDTA for experiments at pH 7.2 and HEPES was replaced by3 mM MES for experiments were TRPV1 was activated by pH 6. The capsaicin stock was prepared at 10 mM in absolute ethanol and then diluted to 250 nM in recording solution. Experiments were performed at room temperature (24 °C) and mean current values were measured after channel activation had reached the steady-state. Pipettes were fabricated with borosilicate glass and had a resistance of 5 MΩ.

Currents were obtained using voltage protocols where the holding potential was 0 mV and then stepped from −120 to 120 mV for 100 ms using a square voltage protocol. Briefly, leak currents were obtained in the absence of agonists or dapagliflozin and then excised membrane patches containing TRPV1 channels were exposed to DAPA, capsaicin, capsaicin + DAPA, pH 6 or pH 6 + DAPA. All currents were leak-subtracted and normalized to currents activated either by capsaicin or pH alone.

### 3.5 Immunocytochemistry (ICC)

Vascular smooth muscle cells were isolated from whole mesenteric and renal arteries from Wister rats. Arteries were placed in a smooth muscle dissociation solution (SMDS) containing (mmol·L^−1^); 120 NaCl, 6 KCl, 12 glucose, 10 HEPES and 1.2 MgCl_2_ supplemented with 1.75 mg·ml−1 Collagenase Type IA, 0.9 mg·ml^−1^ protease, 1 mg·ml^−1^ trypsin inhibitor and 1 mg·ml^−1^ bovine serum albumin (Sigma, UK) at 37°C for 45 minutes. Vessels were washed in SMDS and underwent mechanical trituration using a glass Pasteur pipette to liberate vascular smooth muscle cells. The subsequent cell suspension was plated onto 13mm coverslips and left at room temperature (RT, 20-22°C) for 45 minutes to adhere. The VSMCs were fixed with 4% paraformaldehyde for 15 minutes and then stored in a 24-well plate with filtered PBS at 4°C until used for ICC experiments.

The VSMCs were incubated in WGA Texas Red membrane stain (Thermos Fisher Scientific, Invitrogen) for 10 minutes. Cells were washed with PBS and incubated in 100 mmol·L−1 glycine in PBS for 5 minutes and subsequently in blocking solution (PBS containing 0.1% Triton X-100 and 10% FBS in PBS) for 45 minutes. Cells then underwent an overnight incubation with a mouse anti-SGLT2 antibody (dilution 1:200, Santa Cruz, Texas, USA), then washed in PBS and incubated with a donkey anti-mouse secondary antibody conjugated to Alexa Fluor 488 (dilution 1:100, Thermo-Fisher, Paisley, UK). The coverslips were washed in PBS before being mounted onto a microscope slide with 4’,6-diamidino-2-phenylindole (DAPI) (Vectashield, Vector Laboratories, Burlingame, CA) mounting medium.

### 3.6 Immunohistochemistry

After completion of functional myography experiments, mesenteric and renal arterial segments were fixed *in situ* in myograph chambers with 4% paraformaldehyde (J61899, Thermo Scientific) for 1 hour at room temperature. Fixed arteries were washed with PBS, removed from wire myograph chambers by cutting open laterally and then separated into 2 fixed segments. Arteries were blocked for 90 minutes at room temperature with blocking buffer (1% BSA, 0.5% Triton X-100, 0.05% Tween 20 in PBS) and incubated overnight at 4°C with guinea pig anti-TRPV1 (Ab10295, 1:1000, Abcam) and goat anti-CGRP (Ab36001, 1:1000, Abcam), rabbit anti-NHE1 (PA5115917, 1:1000, Invitrogen), mouse anti-SGLT2 (sc-393350, 1:200, Santa Cruz), rabbit anti-smooth muscle Myosin heavy chain 11 (ab125884, 1:500, Abcam) and Goat anti mouse CD31/PECAM-1 (AF3628-SP, 1:150, R&D Systems) diluted in blocking buffer. This was followed by a secondary antibody incubation with either goat anti-guinea pig (Alexa fluor 488, A11073, Life Technologies), donkey anti-goat (Alexa fluor 633, A21082, Life Technologies), donkey anti-rabbit (Alexa fluor 488, A21206, Life Technologies) or donkey anti-mouse (Alexa fluor 594, A21203, Invitrogen) diluted in blocking buffer for 90 minutes at room temperature. Arteries were then placed in mounting medium (Vectasheild plus Antifade, Vector Laboratories) and laid flat between 2 glass coverslips. Arteries were excited at 405, 488, 536 and 635 nm and fluorescence acquired through a water immersion objective (1.15, NAI, 1024 x 1024 pixels, x40 lens objective, Olympus) using a Fv1000 laser scanning confocal microscope (Olympus, Southend-on-Sea, UK). Z stacks were taken through every arterial segment starting from the arterial wall in 1µM increments using Fluaview version 4.1 and Imaris version 8.0.2 Bitplane software.

### 3.7 Western Blot

Whole protein lysates from mesenteric and renal arteries were prepared using Triton Buffer (Fisher Scientific) supplemented with protease and phosphatase inhibitors (cOmplete, mini and PhosSTOP from Roche). Protein concentrations were determined via the Pierce™ BCA Protein Assay Kit (Thermo Fisher Scientific, Loughborough, UK). The samples were run under reducing conditions with 4-12% Bolt™ Bis-Tris Plus pre-cast gels (Invitrogen) and proteins were transferred to a nitrocellulose membrane. Membranes were blocked for at least 0.5 hour in 3% BSA-PBS and incubated overnight at 4°C with the primary SGLT2 mouse monoclonal antibody (D-6, sc-393350, 1/200 dilution, Santa Cruz). The membranes were incubated with highly adsorbed horseradish peroxidase-conjugated goat anti-mouse IgG (A16078, Fisher Scientific) for 1h at RT and developed using Immobilon™ Western Chemiluminescent HRP Substrate (Millipore). Detection and quantification of chemiluminescence intensities were quantified by using Chemidoc^TM^ imaging system and Image Lab 5.2.1 software (BioRad). Blots were stripped with restore buffer (Fisher Scientific) according to the manufacturer’s instructions and re-probed with a rabbit-anti-actin antibody (#4970, Cell Signalling Technology) and highly-adsorbed horseradish peroxidase-conjugated donkey-anti rabbit IgG (Merks) as above.

### 3.8 Myography

Mesenteric and renal arteries were cut into ∼2mm segments and mounted on 40µm stainless steel wires in a myograph (DMT, Aarhus, Denmark). The myograph chambers contained PSS that was bubbled with 95% oxygen and 5% carbon dioxide at 37°C. Tension in each segment was recorded using LabChart Pro Software (ADInstruments, Oxford, UK).

All vessels were subject to a normalisation procedure (Mulvany and Halpern 1977) to standardise the experimental conditions and arteries were set to an internal circumference 90% of the diameter at in-vivo transmural pressure of 100mmHg (13.3 kPa). Once basal tone was established the VSMC viability was determined by replacing the chambers PSS solution with PSS containing 60mM KCl with a concomitant reduction in NaCl until a sustained contraction was seen. Vessels were then constricted with 10µM of the α1-adrenoreceptor agonist, methoxamine and allowed to plateau before recording the vasorelaxation to 10µM carbachol to determine the endothelial cell integrity.

Arterial segments were preconstricted with 10 µM methoxamine and responses to the SGLT2 inhibitors dapagliflozin, empagliflozin, ertugliflozin (1 to 100µM), the SGLT1 inhibitor mizagliflozin (1-30µM), NHE1 inhibitor cariporide, CGRP (10pM-10nM) and capsaicin (10 µM) determined. Vessels were pre-incubated in the presence or absence of a combination of solvent control dimethyl sulphoxide (DMSO) or pharmacological agents: Linopirdine (pan Kv7 channel blocker, 10µM), BIBN-4096 (CGRP receptor blocker, 1µM), Capsaicin (TRPV1 agonist, 10µM), AMG-517 (TRPV1 blocker, 1µM), AM0902 (TRPA1 channel blocker, 1µM), HMR-1556 (Kv7.1 channel blocker, 10µM), Iberiotoxin (BK_Ca_ channel blocker, 100nM), 4-aminopirydine (4-AP; 1mM), tetraethylammonium (TEA; 1mM) and glibenclamide (K_ATP_ channel blocker, 100nM). To deplete sensory nerves relaxed arteries were challenged with 10µM capsaicin for 5 mins followed by extensive washout over 10 mins. This was repeated twice before contraction with methoxamine. A similar protocol was used for the cariporide pretreatment experiments, except cariporide was left in the bath at the time of contraction.

### 3.9 Reverse transcription quantitative polymerase chain reaction (RT-PCR)

mRNA was extracted from mesenteric and renal artery whole cell lysates using the Monarch Total RNA Miniprep Kit’ (New England Biolabs, Ipswich, Massachusetts, USA). The concentration and purity of the RNA samples were calculated using a NanoDrop 3300 Fluorospectrometer with absorbance set to 260nm and 280nm. Samples were reversed transcribed using the LunaScript RT SuperMix Kit (New England BioLabs, Ipswich, Massachusetts, USA) and relative gene expression was quantified via CFX-96 Real-Time PCR Detection System (RRID:SCR_018064, BioRad, Hertfordshire, UK). Samples were run in duplicate in a BrightWhite 96 well qPCR plate (primer design, Camberley, UK). Each well contained 10ng of cDNA, 300nmol-L of a gene specific primer (thermofisher scientific, USA) and 10uL of Syber green qPCR mastermix (New England Biolabs, Ipswich, Massachusetts, USA) each used as per manufacturer’s instructions. For each gene a no template control (NTC) and NRT sample was run to detect contamination and primer dimers. The quantification cycle (Cq) for each sequence specific primer was determined by a Bio-rad CFX96 Manager 3.0 software and normalised to the housekeeper genes GAPDH and Cytochrome C1 (CYC1), which was expressed as 2^−ΔCq^ value. Table of primers is shown below.

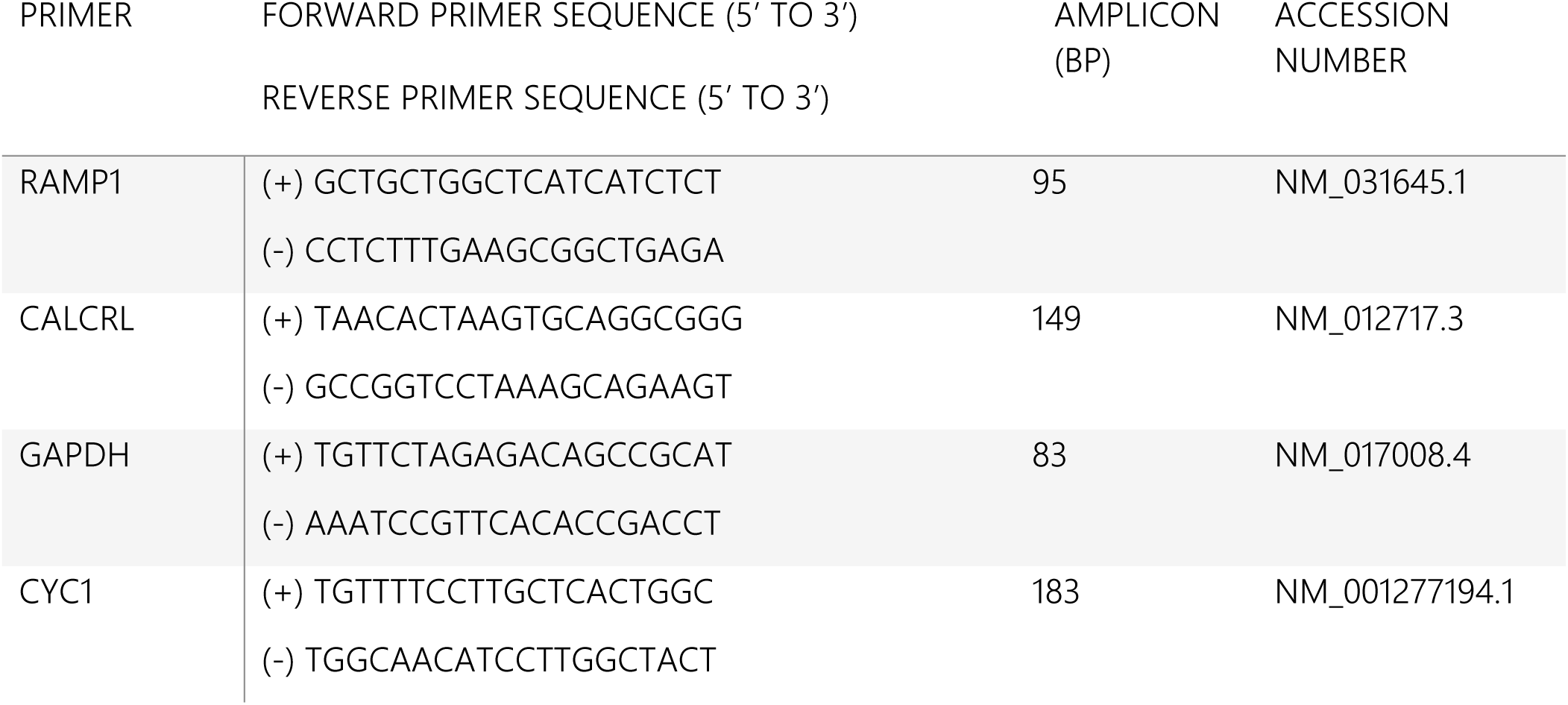

### 3.10 Data and statistical analysis

All values from functional experiments are expressed as mean ± standard error of the mean (SEM) with no less than 5 individual data points, each representing a biological repeat. Measurements of total cell florescence during immunocytochemistry involved five biological repeats with a minimum of five cells to be recorded per sample. For quantification of SGLT2 protein via western blot, a minimum of 3 biological repeats were obtained.

For functional experiments, cumulative concentration effect curves were produced, whereby the contraction produced by 10µM methoxamine at stable tone was taken as the maximal contraction of 100%. The tone of the artery was recorded after each subsequent addition of the pharmacological agent and the values were formulated as a percentage of the maximum contraction. Using GraphPad Prism (**<u>RRID:SCR_002798</u>**, Version 9.0.0) a transformed data set of mean values was generated using X = Log(X), to reduce representative skew. A four parametric linier regression analysis was then performed to produce a concentration effect curve on a log(x) graph with the standard error of mean (SEM).

When comparing multiple groups, a two-way ANOVA was performed followed by a post-hoc Bonferonni or Dunnett’s test. For data comparing 2 groups, an unpaired parametric t-test was performed. Significance values are represented as *P<0.01, **P<0.001, ***P<0.0001 and data sets subject to statistical analysis contained at least 5 animals per group, where N = number of independent values.

## 4 Results

### 4.1 Classifying the expression of SGLT2 in mesenteric and renal arteries from male Wister rats

The expression of SGLT2 within the vasculature is ill-defined, hence we wanted to confirm its expression in different vascular beds. Immunocytochemistry and western blot data demonstrated SGLT2 was expressed in mesenteric and renal arteries from male Wister rats to a similar degree (Fig 1).

**Figure 1.**
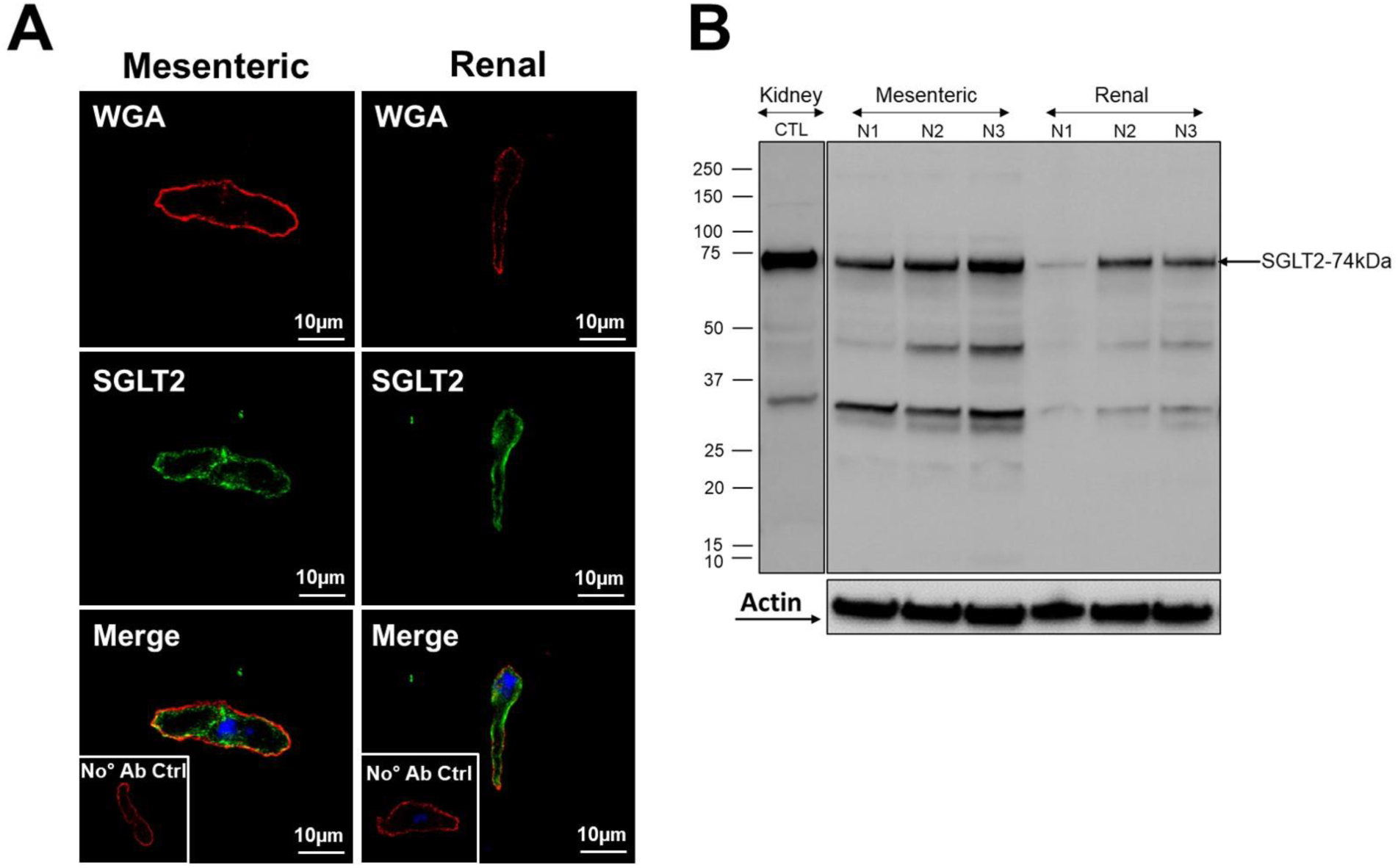
SGLT2 expression in mesenteric and renal arteries. (A) Representative staining of SGLT2 in isolated mesenteric and renal vascular smooth muscle cells (N=5, n=25). (B) Western blot quantification of SGLT2 protein in mesenteric and renal arteries. Whole kidney ran as a positive control (N=3).

### 4.2 SGLT2 inhibitors relax mesenteric arteries in an endothelium independent manner

In 2^nd^ order rat mesenteric artery segments, we determined the vasodilatory effect of 3 structurally different SGLT2 inhibitors (dapagliflozin, empagliflozin and ertugliflozin) and a SGLT1 inhibitor (mizagliflozin). Cumulative application (1µM-100µM) of dapagliflozin, ertugliflozin, empagliflozin and mizagliflozin produced concentration dependent relaxations, with approximate IC_50_ values of 9.5 µM, 7.3 µM, 9.5 µM and 5.52 µM, respectively (Figure 2A, B; n=5-6). The relaxation elicited by each SGLT inhibitor was not affected by the functionality of the endothelium, as indicated by response to carbachol (S1). In contrast to their effect on mesenteric arteries, dapagliflozin and empagliflozin were less effective relaxants of precontracted renal arteries (Fig 2C&D). The NHE1 inhibitor cariporide (Uthman et al. 2018) also relaxed mesenteric arteries (Fig 2E&F) but was ineffective at relaxing pre-contracted renal arteries (Fig 2F).

**Figure 2.**
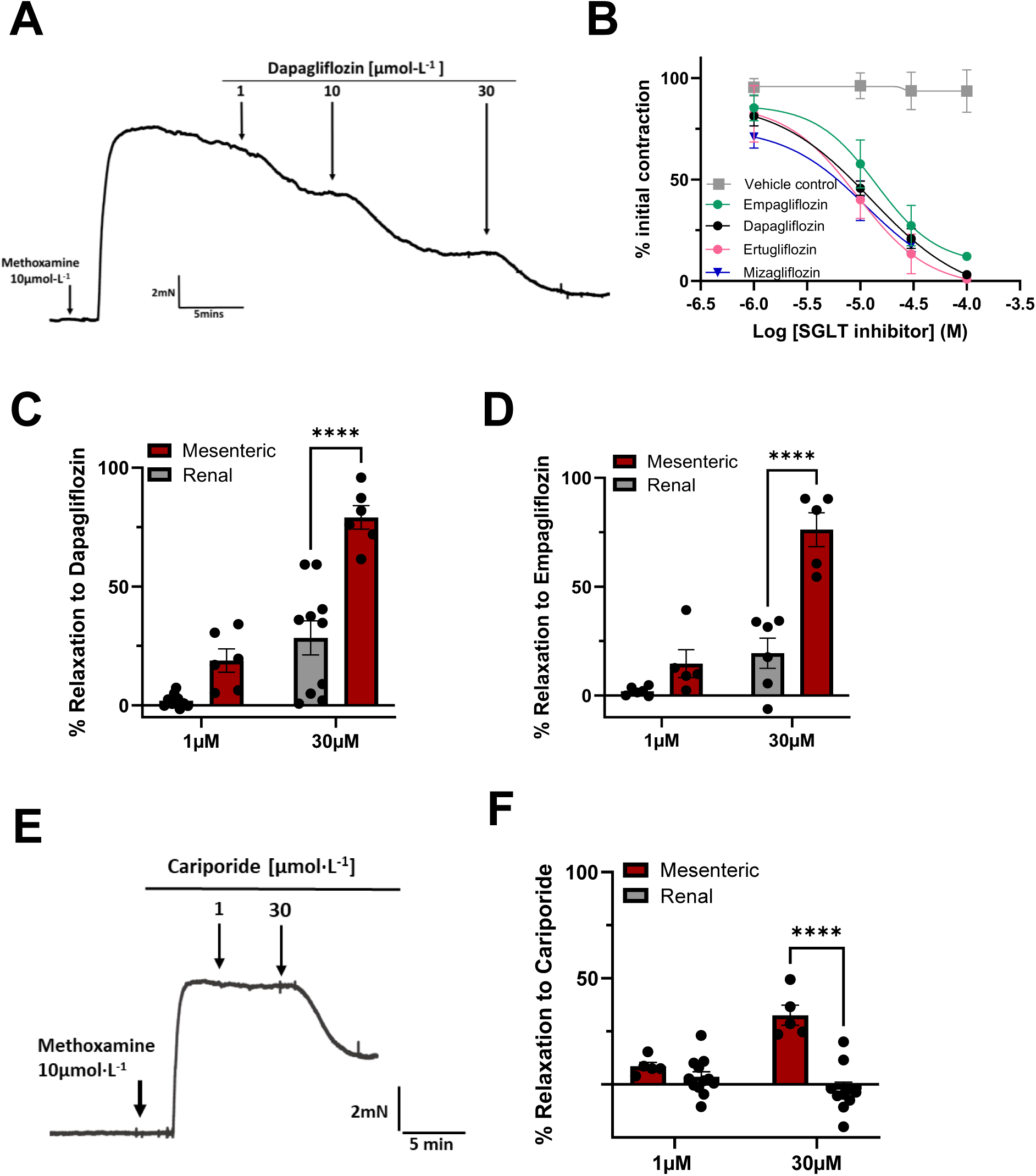
The effect of SGLT2 and SGLT1 inhibitors and NHE1 inhibitor, cariporide on mesenteric and renal artery tone. A shows a representative trace of the effect of dapagliflozin in mesenteric arteries pre-contracted with 10µM methoxamine. B, mean effect of dapagliflozin (black), empagliflozin (green), ertugliflozin (pink) and mizagliflozin (blue), with mean vehicle control in grey, N=5-6. Panel D and E shows the relaxation to dapagliflozin and empagliflozin in renal (grey) compared to mesenteric (red) arteries. The representative trace of the effect of Cariporide (1µM-30µM) in mesenteric arteries is shown in panel E with the mean response to cariporide in mesenteric (red) and renal arteries (grey) shown in panel F. All values are shown as mean ± SEM denoted by the error bars and a 2-way ANOVA with a post hoc Sidack test was used to calculate significance values where ****P<0.0001.

### 4.3 Role of Kv7 channels in SGLT2 inhibitor-induced relaxations

Previous studies implicated voltage-gated potassium channels encoded by *KCNQ* genes (Kv7 channels) in the relaxant response to different SGLT2 inhibitors (Hasan and Hasan 2021; Hasan, Menon, et al. 2022; Hasan, Zerin, et al. 2022). We performed electrophysiological experiments on oocytes co-expressing KCNQ4/KCNQ5 and KCNE4 – a molecular combination found in most arterial smooth muscle (Barrese et al., 2018). Neither dapagliflozin nor empagliflozin at 100 µM had any effect on the current amplitude nor the voltage-dependence of activation of the voltage-dependent potassium currents and neither agent affected the resting membrane potential (Fig 3A-F). However, in mesenteric arteries the relaxation produced by dapagliflozin, empagliflozin and ertugliflozin was significantly attenuated by pre-incubation with the pan-Kv7 channel inhibitor linopirdine (10µM), when compared to DMSO control (Fig 3G and H; n=5-6). Specific blockers of Kv7.1 (HMR1556), BK_Ca_ (iberiotoxin), KATP (glibenclamide) and the non-selective K channel blockers 4-AP and TEA had no effect on relaxations produced by SGLT2 inhibitors (Figure 3I and J). As Kv7 channels are not directly activated by SGLT2 inhibitors we speculated that these channels were recruited by CGRP released from the sensory nerves like other Gs-linked receptor agonists (Stott et al.,2018; Van der Horst et al., 2020). Figure 3K shows that linopirdine inhibited the response to exogenous CGRP to a similar degree as SGLT2 inhibitor-induced relaxations. Consequently, Kv7 channel blockers impaired the relaxant effect of SGLT2 inhibitors by inhibiting the CGRP released by these agents.

**Figure 3.**
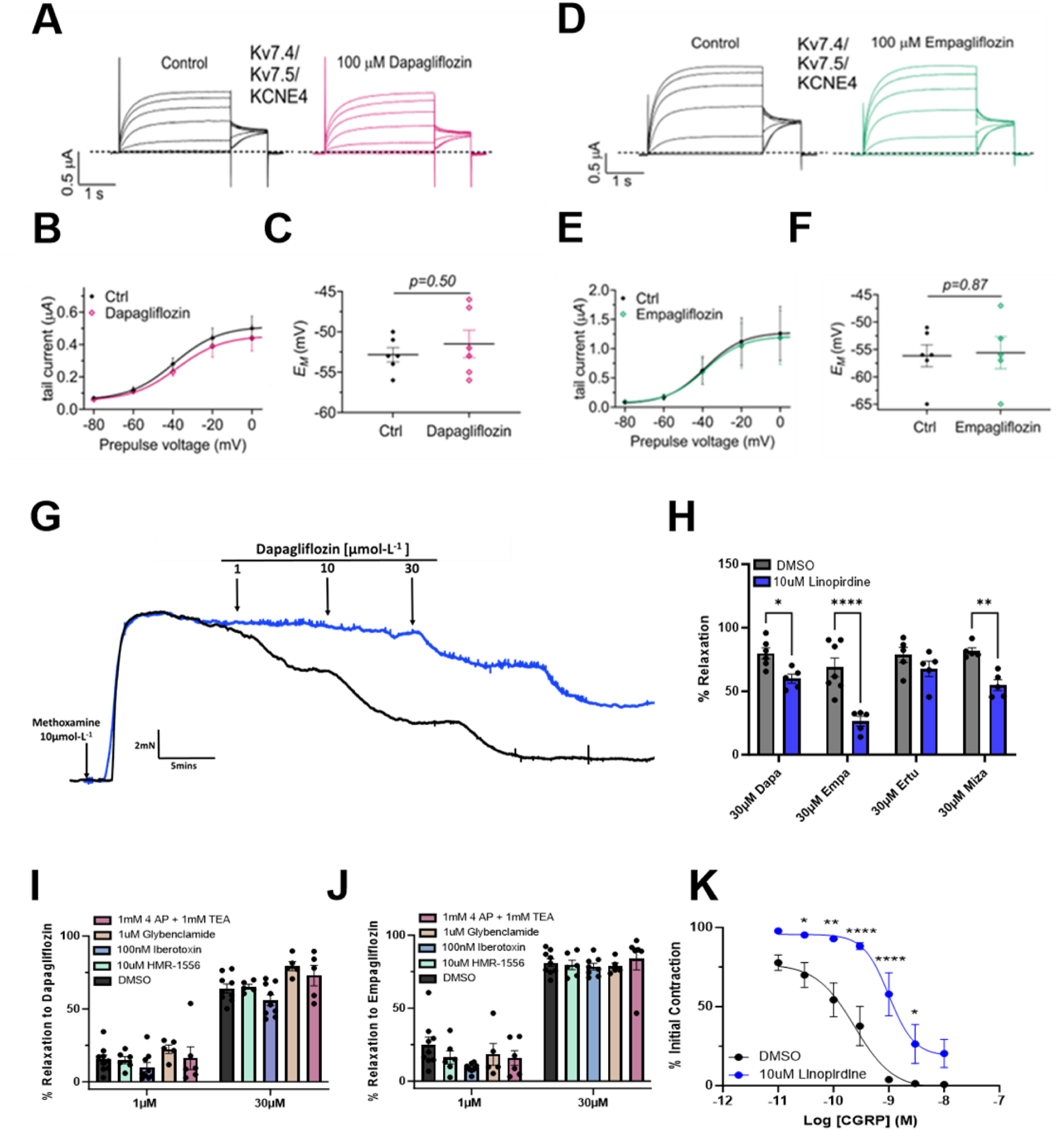
SGLT2 inhibitors and Kv7 channels. Panels A-C shows currents produced by the co-expression of Kv7.4, Kv7.5 and KCNE4 in the absence and presence of 100 µM dapagliflozin. Representative traces in A, mean current-voltage relationship in B and mean membrane potential in C. The effect of 100 µM empagliflozin on currents produced by co-expression of Kv7.4, 7.5 and KCNE4 is shown in panels D-F. Representative traces in D, mean current-voltage relationship in E and mean membrane potential in F. Data are the mean of N oocytes with error bars denoting the SD. Panel G, representative trace of the effect of dapagliflozin on precontracted mesenteric arteries in the presence (Blue) and absence (black) of 10 µM linopirdine. H shows the mean data for relaxations to dapagliflozin, empagliflozin, ertugliflozin and mizagliflozin (all 30 µM) in solvent control (black) and when pre-incubated with 10 µM linopirdine (blue)(N=5-6). All values are expressed as mean ± SEM denoted by the error bars. A two-way statistical ANOVA with a post-hoc Sidak test was used to generate significant values (*=P<0.05, **=P<0.01, ***=P<0.001) (N= number of animals used). The effect of dapagliflozin (I) and empagliflozin (J) in the presence and absence of HMR1556 (green), Iberiotoxin (blue), 4-aminopyridine (AP), tetraethylammonium (TEA, red) and glibenclamide (orange). All data values are shown as mean ±SEM, (N=5-6). K shows the mean relaxation to CGRP in the absence (black) and presence of 10 µM Linopirdine (A-Blue, N=5-8). All values are expressed as mean ± SEM. A two-way statistical ANOVA with a post-hoc bonferoni test was used to generate significant values (*=P<0.05, **=P<0.01, ***=P<0.001, ****=P<0.0001).

### 4.4 Sensory nerve contribution in mesenteric and renal arteries

We speculated that the different effect of the SGLT2 inhibitors between mesenteric and renal arcades was due to a differential abundance of sensory nerves. CGRP is a potent vasodilator that is released by sensory neurones upon TRPV1 channel activation (Aalkjaer et al. 2021). We performed immunohistochemistry experiments with well validated CGRP and TRPV1 antibodies to delineate sensory nerves in both arteries. Figure 4A shows robust TRPV1 and CGRP staining was observed in the adventitia of mesenteric arteries with comparatively little staining in the smooth muscle or endothelial layers (Figure S2). In contrast, negligible TRPV1 or CGRP staining was identified in the adventitial, medial or inner layers of the renal artery (Fig 4B and Fig S2). Thus, mesenteric arteries exhibit robust sensory nerve networks that are not present in renal arteries. In line with this observation, application of the TRPV1 activator capsaicin produced a BIBN-4096-sensitive relaxation of pre-contracted mesenteric arteries but had no effect in precontracted renal arteries (Fig 4C). Figure 4D shows that cumulative application of CGRP (10pmol·L^-1^–10nmol·L^-1^) produced a maximal relaxation of 99.3±0.43% in pre-contracted mesenteric arteries, which was significantly attenuated by BIBN-4096 (n=5-8). Strikingly, CGRP failed to relax renal arteries (Fig 4D). Quantitative PCR revealed no difference in the transcript abundance of *Calrlr* between the two vascular beds, however *Ramp1* was expressed at significantly lower levels in renal compared to mesenteric arteries (Fig 4E). These experiments show that CGRP can relax preconstricted mesenteric arteries when applied exogenously or from endogenous release by activating TRPV1 channels on perivascular sensory nerves. Thus, the reduced relaxation to dapagliflozin and empagliflozin in renal arteries could be underpinned by an absence of sensory nerves and reduced *Ramp1* subunit expression.

**Figure 4.**
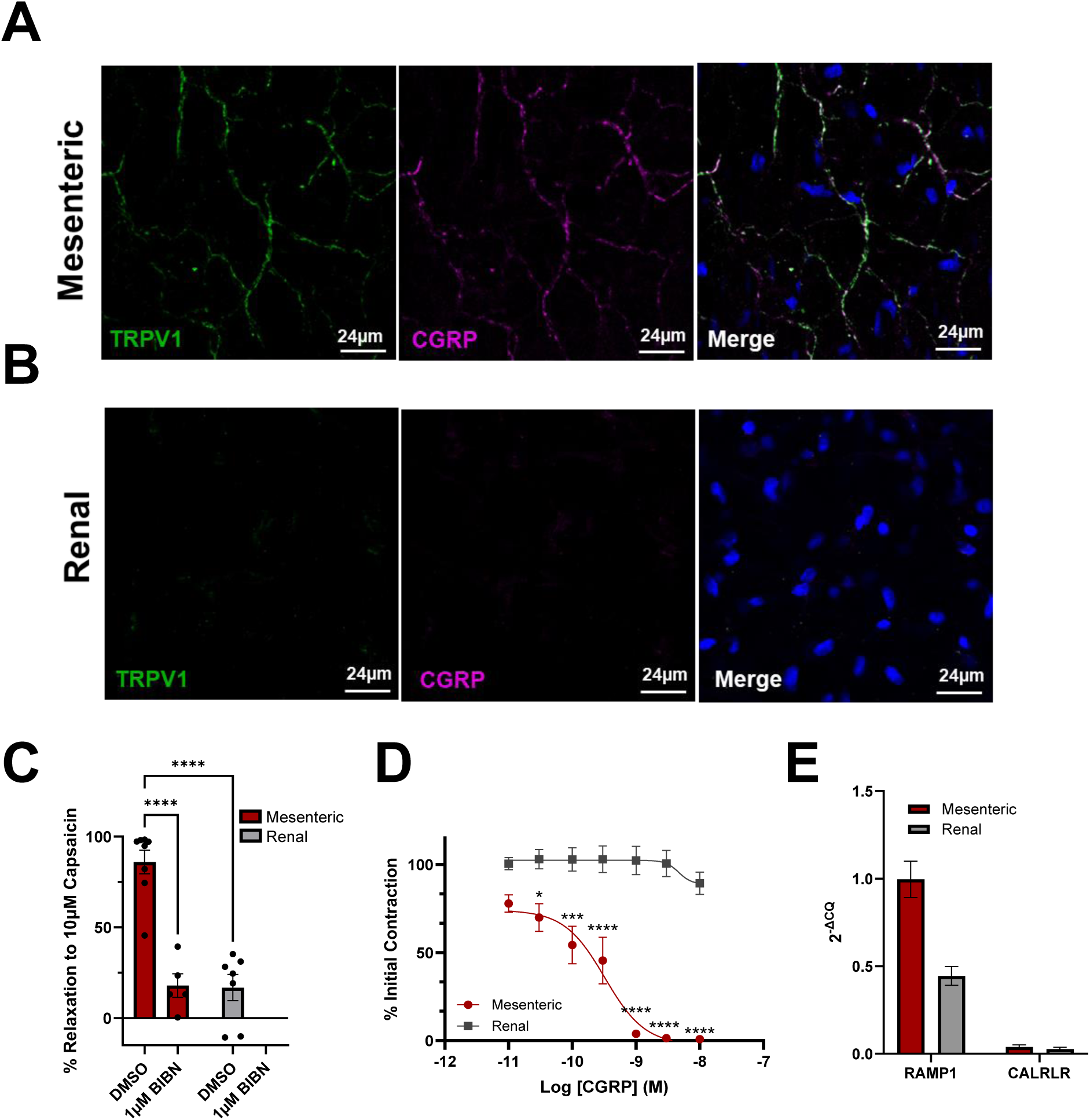
Mesenteric arteries have a greater sensory nerve contribution and response to CGRP and Capsaicin compared to renal arteries. Panels A and B show representative labelling for TRPV1 (green) and CGRP (magenta) indicative of sensory nerve presence in the adventitia of whole mesenteric (A, N=3) and renal arteries (B, N=4). Nuclei were labelled in blue. C shows the percentage relaxation to 10µM Capsaicin in mesenteric (red) and renal (grey) arteries in the presence and absence of 1µM BIBN, (N=6-8). D shows the relaxation to CGRP in mesenteric (red) and renal (grey) arteries, (N=5-8). E is the mean relative transcript abundance of RAMP1 and CALRLR in mesenteric (red) and renal (grey) arteries (E, N=6). All data is represented as mean ±SEM denoted by the error bars. A two-way statistical ANOVA with a post-hoc Sidak test was used to generate significant values (*=P<0.05, **=P<0.01, ***=P<0.001, ****=P<0.0001) (N= number of animals used).

### 4.5 Dapagliflozin and empagliflozin evoked relaxations are sensitive to CGRP blockade

To identify the role for CGRP in SGLT2 inhibitor induced relaxations, we applied dapagliflozin, empagliflozin and cariporide to mesenteric arteries pre-incubated with either DMSO (control), or BIBN-4096 (1 µM; Fig 5A-B; n= 6). The relaxation to both dapagliflozin and empagliflozin was significantly attenuated by 1 µM BIBN-4096 although the attenuation produced by BIBN-4096 was 2-fold greater for empagliflozin compared to dapagliflozin induced relaxations. Cariporide-induced relaxations of mesenteric artery were also markedly impaired by BIBN-4096 pretreatment (Fig 5C). Hence, relaxations evoked by SGLT2 inhibitors and the NHE1 blocker were sensitive to CGRP receptor blockade.

**Figure 5.**
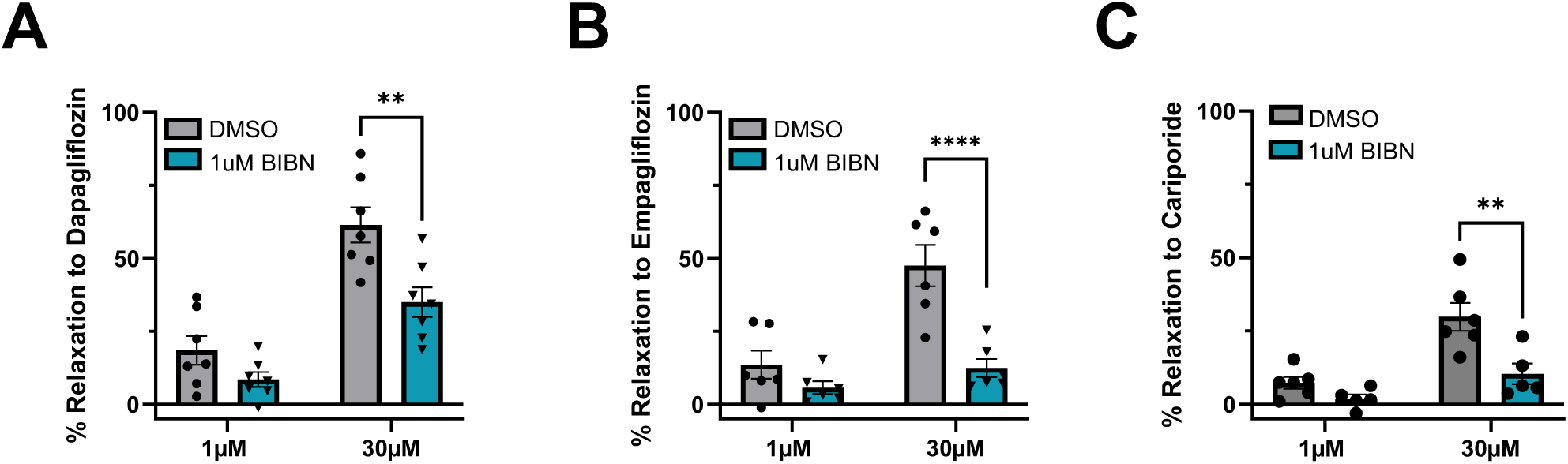
Dapagliflozin, empagliflozin and Cariporide induced relaxations are blocked by CGRP receptor antagonist. The mean effect of dapagliflozin (A, N=6), empagliflozin (B, N=7) and Cariporide (C, N=5) on precontracted mesenteric arteries in the presence of DMSO (solvent control, grey), 1 µM BIBN (teal). All values are expressed as mean ± SEM. A two-way statistical ANOVA with a post-hoc Sidak test was used to generate significant values (*=P<0.05, **=P<0.01, ***=P<0.001) (N= number of animals used).

### 4.6 Dapagliflozin and empagliflozin induced relaxations are sensitive to TRPV1 but not TRPA1 blockade

We hypothesised that SGLT2 inhibitors relaxed mesenteric arteries through provoking CGRP release from sensory nerves. To confirm this, we took two approaches – depletion of CGRP stores through treatment with three 5 min applications of capsaicin (10µM) followed by wash out of the bathing solution and direct block of TRPV1 with AMG-517 (1µM). Both manoeuvres abrogated the relaxations produced by 1 µM capsaicin (Figure 6A-B). Figure 6C shows that the relaxation to cariporide was also diminished in the presence of the TRPV1 blocker AMG-517. The relaxations produced by both dapagliflozin and empagliflozin were significantly attenuated after treatment with capsaicin (Fig 6D & E, N=6-10). The TRPV1 blocker, AMG-517 attenuated the dapagliflozin relaxation to the same extent as capsaicin pretreatment, whereas the empagliflozin relaxation was further attenuated from a ∼60% relaxation to ∼13% (Fig 6D-E). The relaxation to dapagliflozin was not affected when mesenteric arteries were pre-incubated with the TRPA1 blocker AM0902 (1µM, Fig 6F). None of the pharmacological modulators affected methoxamine-induced contractions (Fig 6G). Hence, relaxations induced by SGLT2 inhibitors are dependent upon activation of the TRPV1 channel on perivascular sensory nerves and subsequent CGRP release.

**Figure 6.**
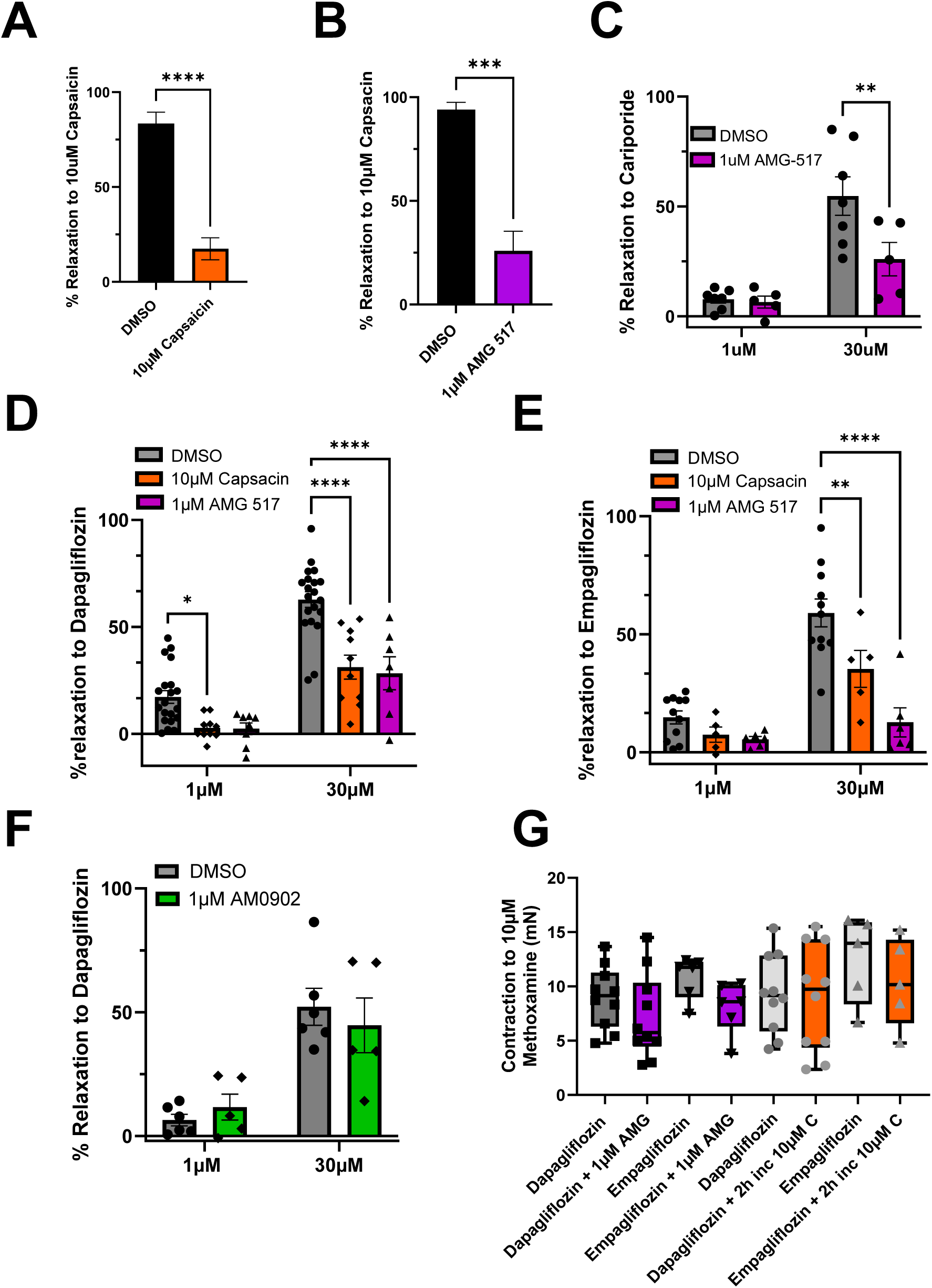
Dapagliflozin and Empagliflozin relaxations are impaired by TRPV1 blocker AMG −517 and Capsaicin. Panels A and B show the mean effect of 10µM capsaicin in arteries after sensory nerve depletion (A, 3 −5 min pulses of capsaicin then wash out, N=6) or in the presence of the TRPV1 blocker AMG −517 (B, N=5). Mean relaxations produced by 1µM and 30µM cariporide (C, N=5-7) empagliflozin (D, N=6-10) and dapagliflozin (E, N=7-10) in mesenteric arteries pre-incubated in 1 µM AMG-517 (purple) or after sensory nerve depletion with 10µM Capsaicin (orange). F shows the mean response to 1µM and 30µM dapagliflozin in the presence (green) and absence (grey) of the TRPA1 blocker AM0902 (N=5). G shows the mean amplitude of the contraction produced by 10µM Methoxamine under the different conditions. All values are expressed as mean ± SEM. A two-way statistical ANOVA with a post-hoc Sidak or bonferroni test was used to generate significant values (*\*\*=P<0.01, ***=P<0.001, ****=P<0.0001*). (*N*=) number of animals used.

### 4.7 Effect of SGLT2 inhibitors on heterologously expressed TRPV1 currents

To determine whether dapagliflozin could directly activate TRPV1 channels, we performed patch-clamp experiments using outside-out excised membrane patches of HEK293 cells expressing rat TRPV1 (rTRPV1). As shown in Figure 7, we first obtained the leak currents (in the absence of agonist, grey traces) elicited by a square voltage pulse to −120 mV followed by a pulse to 120 mV, then used the same voltage protocol to assess currents after exposing the patches to either 30 μM or 100 μM dapagliflozin for 5 mins and, finally, to 250 nM capsaicin alone (black traces). All currents were leak-subtracted and normalized to the current obtained at +120 mV (Fig 7C). Currents after exposure to 30 μM dapagliflozin were 12.9±2.2% and 7.6±3.7% after 100 μM dapagliflozin of the currents elicited by the TRPV1 agonist, capsaicin (Figure 7C, n=6). These data indicate that TRPV1 is not directly activated by dapagliflozin.

**Figure 7.**
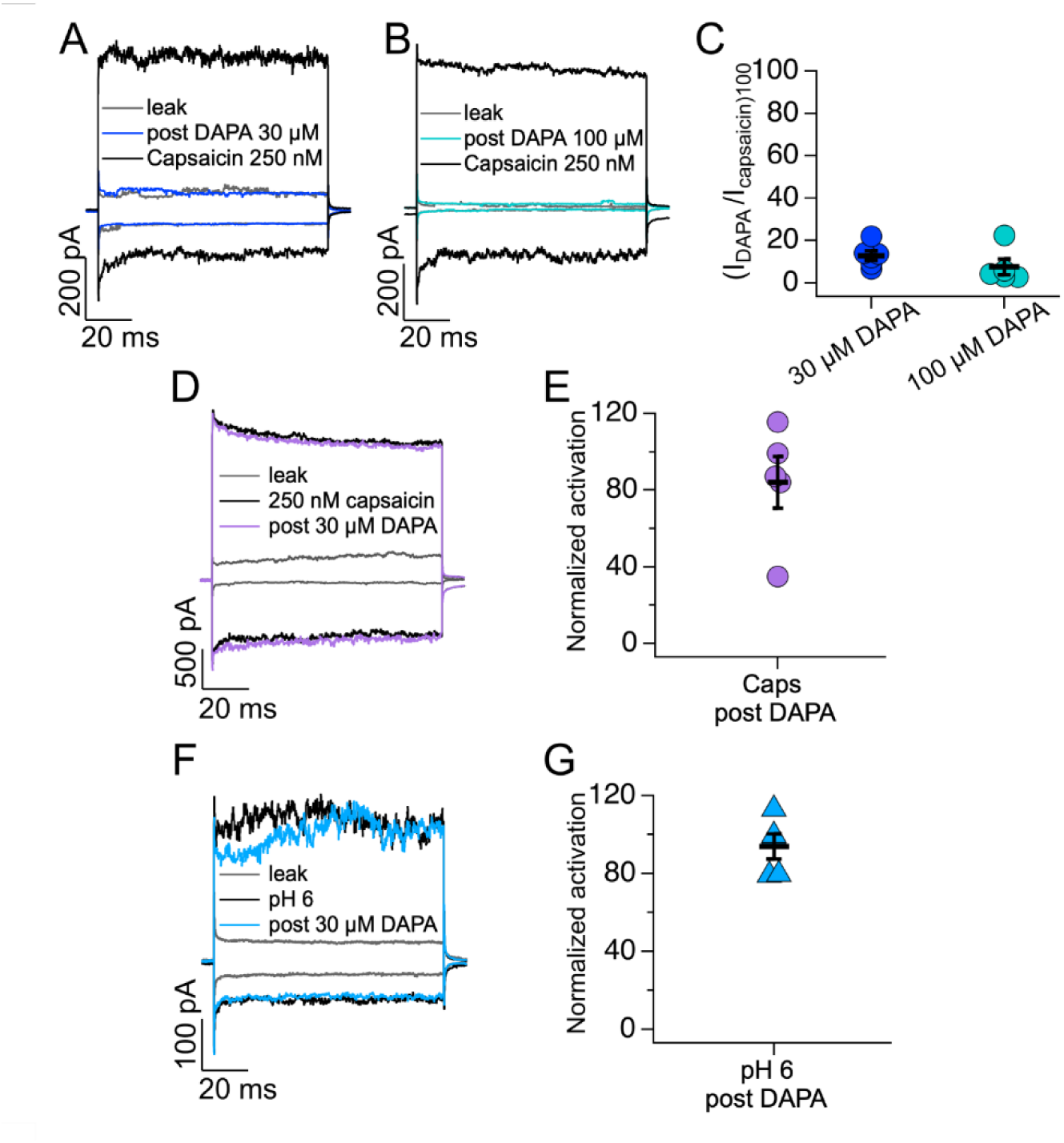
SGLT2 inhibitors do not activate TRPV1 directly. (A and B) Representative traces of currents at +120 and −120 mV from outside-out membrane patches of HEK293 cells expressing TRPV1. Leak currents were obtained in the absence of any agonist (grey) and after 5 min application of dapagliflozin (DAPA) 30 μM (blue traces, panel A) and 100 μM (green traces, panel B). Black trace shows the subsequent effect of 250 nM capsaicin. (C) Average data for experiments in (A and B). Currents were leak-subtracted, and data was normalized to activation by capsaicin 250 nM in the steady-state at +120 mV (N = 6 and N = 5 for DAPA 30 μM and 100 μM, respectively). (D) Representative traces of currents at +120 and −120 mV from outside-out membrane patches of HEK293 cells expressing TRPV1 in control conditions (grey), after application with 250 nM capsaicin (black traces) and after application of 250 nM capsaicin + 30 μM DAPA for 5 mins (lilac traces). (E) The data in (D) was normalized by dividing the currents obtained at +120 mV in response to 250 nM capsaicin + 30 μM DAPA by the currents in response to 250 nM capsaicin alone; (N = 5) (F) Representative traces of currents at +120 and −120 mV from outside-out membrane patches of HEK293 cells expressing TRPV1 under control conditions (grey), after activation of TRPV1 by pH 6 (black) and after 5 min application of 30 μM DAPA to pH 6 conditions (blue). (G) The data in (F) were normalized by dividing the currents obtained at +120 mV in response to pH 6 + 30 μM DAPA by the currents in response to pH 6 alone; (N = 6). Group data are reported as the mean ± standard error of the mean.

Next, we studied whether dapagliflozin could potentiate capsaicin-or low pH -activated TRPV1 currents. For this set of experiments, we first recorded leak currents (grey traces), then activated TRPV1 in outside-out excised membrane patches of HEK293 cells with either a sub saturating concentration (250 nM) of capsaicin or with a solution at pH 6, then washed the membranes patches to close the channels and exposed the patches to 30 μM dapagliflozin for 5 mins and remeasured the currents in the presence of 250 nM capsaicin or solution with low extracellular pH (Figs 7D-G). The results from these experiments indicate that 30 μM dapagliflozin did not potentiate TRPV1 currents activated by 250 nM capsaicin (Fig 7E; 84.1±13.5%, n = 5) or by low pH (Fig 7G 94±6.5%, n =5). Thus, SGLT2 inhibitors neither activate TRPV1 directly nor sensitize the channel to known mediators.

### 4.8 NHE1 colocalises with TRPV1 and cariporide prevents SGLT2 inhibitor-induced responses

Since SGLT2 inhibitors and mizagliflozin are known to block NHE we postulated that SGLT2 inhibitors promoted release of CGRP from sensory nerves by inhibiting NHE and producing a localised pH change sufficient to activate TRPV1. To test this, we stained mesenteric arteries with SGLT2 and NHE1 and used TRPV1 to delineate the sensory nerves. As shown in Figure 8A, prominent NHE1 staining was observed in the adventitia of mesenteric arteries co-localised with TRPV1 and some staining in the smooth muscle and endothelial layers. No NHE staining was observed in the adventitial layers of renal arteries. In contrast, negligible staining of SGLT2 was observed in the adventitial and endothelial layers but traces of staining were identified in the smooth muscle layer of mesenteric arteries (Fig 8B). No staining for NHE1 or SGLT2 was detected in renal artery adventitia (Fig S3). Therefore, NHE1 but not SGLT2 colocalised with TRPV1 in sensory nerves in the mesenteric artery.

**Figure 8.**
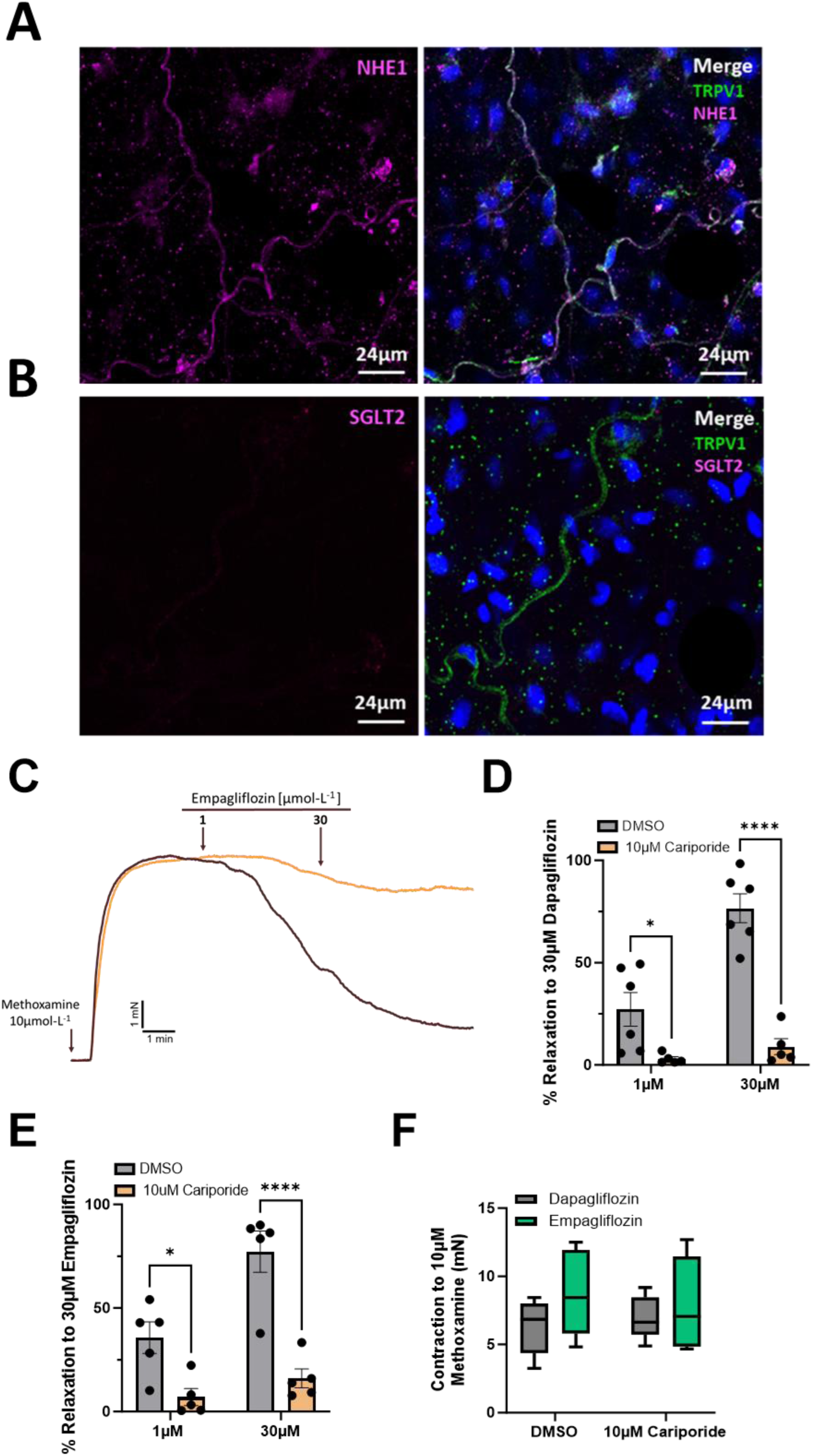
NHE1 colocalises with TRPV1 and cariporide attenuates SHLT2 inhibitor-indued relaxations. Representative immunocytochemistry labelling in the adventitia of whole mesenteric arteries with NHE1 (magenta) and TRPV1 (green) in panel A (N=3) and SGLT2 (magenta) and TRPV1 (green) in panel B (N=3). Nuclei were labelled in blue. A representative trace of the relaxation to empagliflozin in mesenteric arteries pre-contracted with 10µM Methoxamine in the presence (orange) and absence (black) of cariporide (C). The mean data for the response to 1µM and 30µM dapagliflozin (N=5) and empagliflozin (N=5) in the presence (orange) and absence (grey) of 10µM cariporide is shown in Pannel D and E. F shows the mean amplitude of the contraction produced by 10µM Methoxamine under the different conditions. All values are expressed as mean ± SEM. A two-way statistical ANOVA with a post-hoc Sidak test was used to generate significant values (*=P<0.05, **=P<0.01, ***=P<0.001, ****=P<0.0001). (N=) number of animals used.

In mesenteric arteries pretreated with 30µM cariporide, the relaxations to 30µM dapagliflozin or 30µM empagliflozin were significantly smaller (p<0.001) than relaxations produced in the time matched control segments (Fig 8C-E). The amplitude of methoxamine-induced contractions was not significantly different in cariporide-treated arteries compared to DMSO treated (Fig 8F). These experiments support that vasodilatory effects of SGLT2 inhibitors were mediated by NHE1 inhibition.

## 5 Discussion

This study investigated the underlying mechanism of SGLT2 inhibitor-induced vasorelaxation and a possible role for NHE1 inhibition. Our results show that cumulative application of structurally different SGLT2 inhibitors relaxed mesenteric arteries equipotently, in contrast to their ability to block SGLT2 transport for which the Ki for ertugliflozin, dapagliflozin and empagliflozin is 0.88nM<1.1nM<3.1nM respectively (Grempler et al. 2012; Wright 2021). These relaxations were sensitive to TRPV1 and Kv7 channel blockade, however electrophysiology recordings showed that SGLT2 inhibitors did not activate Kv7 and TRPV1 channels directly. In addition, relaxations to dapagliflozin and empagliflozin were attenuated markedly by CGRP receptor blockade and both agents were relatively ineffective in renal arteries. Our data demonstrated this was due to mesenteric arteries being richly innervated by sensory nerves as evinced by staining for CGRP and TRPV1 in the adventitia whereas renal arteries were not. Moreover, renal arteries expressed far less of the CGRP receptor component *Ramp1* and were refractory to exogenous capsaicin application. The NHE blocker cariporide also relaxed mesenteric arteries, which was sensitive to CGRP receptor and TRPV1 blockade, but had no effect in renal arteries. Strikingly, only NHE1 colocalised with TRPV1 in sensory nerves and pre-application of cariporide attenuated the relaxations produced by empagliflozin and dapagliflozin. Our data provide strong evidence that SGLT2 inhibitors influence arterial reactivity by promoting the release of CGRP and potentially other neuropeptides from sensory nerves mediated by the well-known effect of these agents on NHE1 (Baartscheer et al. 2017; Uthman et al. 2018; De Stefano et al. 2021). As the density of sensory nerve innervation varies across the vasculature and within an artery (see staining in Figs 4 & 9) and sensory nerves can modulate endothelium-derived regulators, this seminal finding explains much of the variability in data seen in previous publications (e.g., Hasan and Hasan 2021; Hasan, Menon, et al. 2022; Hasan, Zerin, et al. 2022).

**Figure 9.**
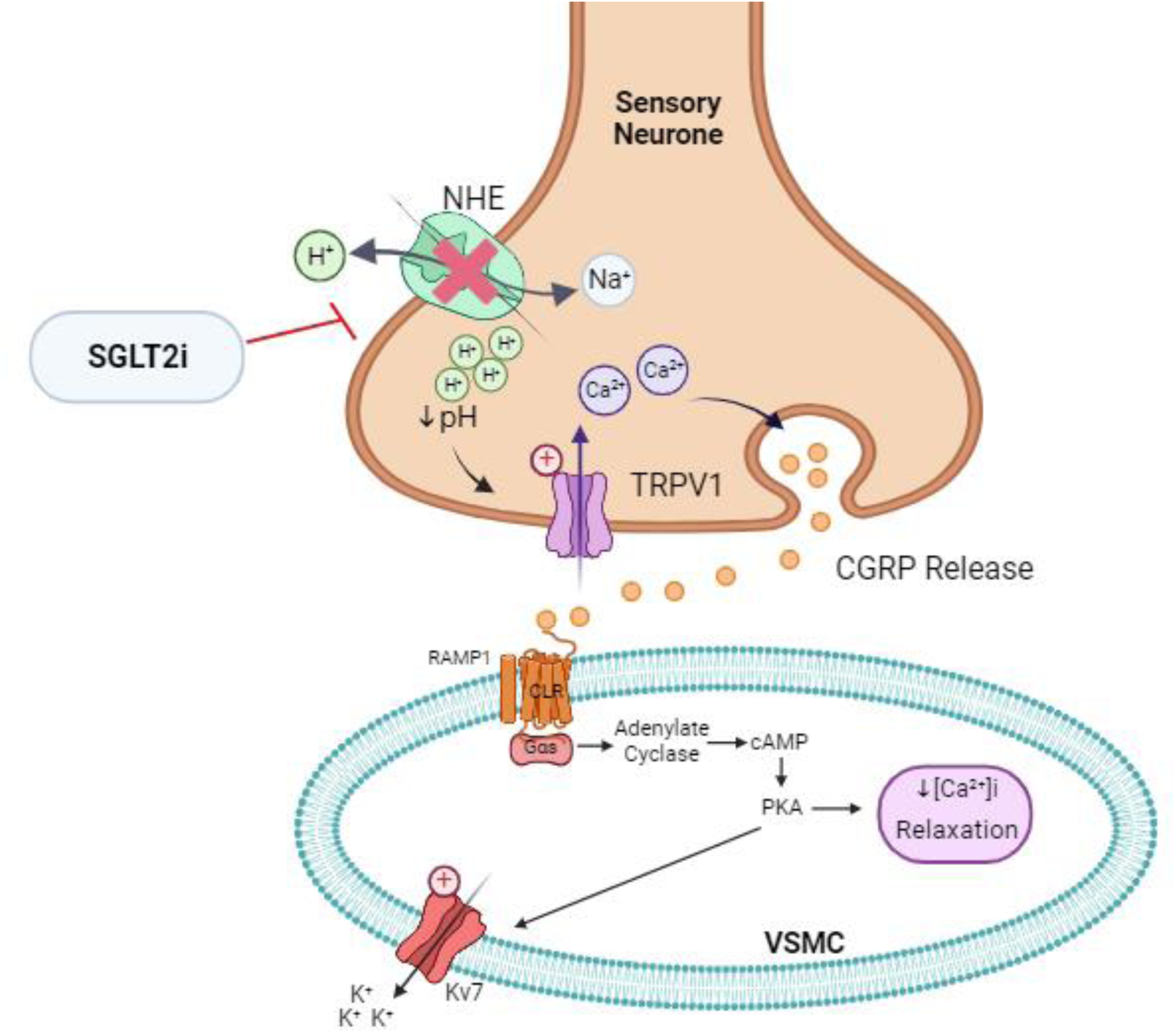
Pathway of SGLT2 induced relaxation in mesenteric arteries. SGLT2 inhibitors acts on the sodium hydrogen exchanger (NHE) to induce vasorelaxation. Hydrogen (H), Sodium (Na), calcium (Ca^2+^), calcitonin-gene related peptide (CGRP), transient receptor potential vanilloid 1 (TRPV1), calcitonin receptor-like receptor (CLR), receptor activity-modifying protein 1 (RAMP1), cyclic guanosine monophosphate (cGMP), protein kinase A (PKA), voltage gated potassium channel (Kv7).

### 5.1 SGLT2 inhibitors induce arterial relaxations

Previous studies showed that empagliflozin, dapagliflozin and canagliflozin relaxed rabbit aortic rings and rat mesenteric arteries, but the underlying mechanisms were ill-defined and often contradictory. In rabbit aortic rings dapagliflozin- and empagliflozin-mediated relaxations were impaired by protein kinase G inhibitors and the non-specific Kv channel blocker 4-aminopyridine (Li et al. 2018; Seo et al. 2020) but specific blockers of Kv subfamilies (Kv1.5, Kv2.1 and Kv7s) did not impair relaxations. In contrast, dapagliflozin, empagliflozin and canagliflozin relaxed mesenteric resistance arteries in an endothelium-independent manner that involved, in part, activation of Kv1.5 and Kv7 potassium channels (Hasan and Hasan 2021; Hasan, Menon, et al. 2022; Hasan, Zerin, et al. 2022) without an effect on protein kinase G although no direct electrophysiology was presented. In line with previous arterial studies, we show that the relaxation produced by SGLT2 inhibitors dapagliflozin, empagliflozin and ertugliflozin and SGLT1 inhibitor mizagliflozin in pre-contracted mesenteric arteries were sensitive to linopirdine and not to other Kv channel blockers (HMR-1556, Iberiotoxin, 4-aminopirydine, tetraethylammonium and glibenclamide). However, neither dapagliflozin nor empagliflozin enhanced potassium currents in oocytes expressing Kv7.4, Kv7.5 and KCNE4 (the combination in arterial smooth muscle, Barrese et al., 2018). Kv7 channel blockers impaired CGRP-induced relaxations in cerebral and mesenteric arteries (Chadha et al. 2014; Stott et al. 2018, present study) so we propose that these channels are a functional endpoint of a relaxant cascade that starts with TRPV1 in the sensory nerves.

### 5.2 The role of CGRP in arterial effects of SGLT2 inhibitors

#### ELIZABETH PLEASE FIX THE FIGURE FOR THE TRPVI CHANNEL – I SUGGEST YOU DON’T MAKE IT BLOCKY BUT USE A CHARACTER LIKE THE KV7 CHANNEL

Our hypothesis (summarised in Fig 9) is that the arterial relaxation produced by the structurally dissimilar SGLT2 and SGLT1 inhibitors was mediated predominantly by CGRP release from perivascular sensory nerves (Aalkjaer et al., 2021) that are lacking in renal arteries. Thus, blocking the CGRP receptor with BIBN-4096 impaired the empagliflozin and dapagliflozin-induced relaxations of mesenteric artery although the lack of complete inhibition suggests other neuropeptides like substance P may also be released. The SGLT2 inhibitor-induced relaxation was equally prevented by application of the TRPV1 blocker AMG-517 and by depletion of CGRP through capsaicin treatment but not by the TRPA1 inhibitor AM0902. This suggests that the sensory nerves in mesenteric arteries contain homomeric TRPV1 channels and not TRPV1/TRPA1 heteromers. Moreover, TRPA1 homomers are unlikely to contribute to SGLT2 inhibitor responses.

TRPV1 is a polymodal cation channel that is regulated by various exogenous and endogenous activators. These include noxious chemicals (capsaicin or vanilloids), low pH (<6.0), high temperatures >43°C (Caterina et al. 1997), lipid mediators (i.e., anandamide), Lipoxygenase products (e.g., LTB4) (Huang et al. 2002) and several signalling molecules (NGF, ATP and PAR-2 agonists) (Chuang et al. 2001; Fernandes et al. 2012). The subsequent influx of cations through TRPV1 is sufficient to promote fusion of synaptic vesicles containing CGRP and other neuropeptides. Studies using TRPV1 knockout mice show activation of TRPV1 leads to vasorelaxation in mesenteric arteries linked to an elevation in PKA (Yang et al. 2010). Our data are consistent with SGLT2 inhibitors relaxing mesenteric arteries through promoting CGRP release in a manner dependent upon TRPV1 channels. However, in over-expression systems dapagliflozin and empagliflozin failed to either activate TRPV1 currents or enhance the effect of low pH or capsaicin, suggesting that these agents do not work directly on the channel. Therefore, TRPV1 activation is a consequence of an additional mechanism.

#### 5.3 NHE1 and arterial relaxation

NHE-1 plays a primary role in cardiomyocytes and vascular smooth muscle cells to maintain cellular pH levels at approximately 7.2 (Karmazyn et al. 1999; Wichaiyo and Saengklub 2022). Altered NHE expression and activity have been linked to severe cardiac events, where during ischemia the pH change activates NHE leading to cardiac injury (Wichaiyo and Saengklub 2022) and much of the cardioprotective effects of SGLT2 inhibitors have been linked to reduced Na^+^ load and pH development in cardiomyocytes (Baartscheer et al. 2017; Uthman et al. 2018; De Stefano et al. 2021). However, an arterial role for NHE1 in the clinical benefit of SGLT2 inhibitors has not been considered. NHE1 activation is linked to vasoconstriction and enhances the myogenic response in mouse resistance arteries (Artamonov et al. 2018). Increased NHE1 activity is also implicated in pulmonary artery hypertension, proliferation and remodelling (Lade et al. 2023), with the protein regulating pH or acting as a protein anchor. We propose that NHE1 located in the sensory nerves also has a profound effect in arteries because they influence TRPV1 activity and the subsequent cation influx precipitates vesicular release of potent vasodilators including CGRP. Thus, in the present study, relaxations to SGLT2 inhibitors and the NHE1 blocker cariporide (Uthman et al. 2018) were prevented by the CGRP receptor antagonist and TRPV1 blocker. Moreover, pre-application of cariporide abrogated the response to dapagliflozin and empagliflozin. Crucially, only NHE1 and not SGLT2 was identified in sensory nerves unlike in the proximal convoluted tubule where SGLT2 and NHE1 cohabit the same microdomain (Pessoa et al. 2014). However, inhibition of NHE1 would lead to intracellular acidification and TRPV1 is activated by external protons so the precise mechanism underlying transmitter release from sensory nerves remains to be determined. Moreover, as the SGLT2 inhibitors produced significantly greater relaxation than cariporide it is possible that another mechanism is also involved.

## 6 Conclusion

CGRP is a potent vasodilator of many vascular beds (Aalkjaer et al. 2021), is a safeguard against cardiac ischemia and promotes cardiac contractility in failing hearts (MaassenVanDenBrink et al. 2016). Our data reveal that the beneficial effects of SGLT2 inhibitors stem from an ability to release cardioprotective CGRP into stressed circulations. The ensuing vasodilatation allied to anti-inflammatory, anti-fibrotic and pro-inotropic actions of CGRP will support effective circulation and help to ameliorate cardiovascular stress.

### 6.1 Perspectives

#### What was known?

SGLT2 inhibitors have a beneficial cardiovascular effect in subjects independent of circulating glucose.

SGLT2 inhibitors can relax arteries ex vivo by an ill-defined mechanism.

#### What does this study add?

This work shows that relaxations to three SGLT2 inhibitors is dependent upon CGRP receptor activation and is reliant upon sensory nerve innervation.

Inhibition of the NHE by the SGLT2 inhibitors is implicated in the vasodilatory effect.

#### How will this study add benefit?

SGLT2 are a recommended front line treatment of heart failure. Understanding the mechanisms that underlie the arterial vasodilatation by these agents will inform future use and treatment strategies.

## FUNDING

EF was supported by a PhD studentship (FS/PhD/21/2912) from the British Heart Foundation awarded to IG. GWA was supported by the US National Institute of General Medical Sciences (GM130377). KER was supported by the US National Institute of Neurological Disorders and Stroke (T32NS045540). TAJ was funded by the Lundbeck Foundation (grant R323-2018-3674) and TR was funded by Dirección General de Asuntos del Personal Académico (DGAPA)-Programa de Apoyo a Proyectos de Investigación e Innovación Tecnológica (PAPIIT) (IN200423).

## ACKNOWLEDGEMENTS

Dr Lillian Wallis at the Department of Pharmacology, Oxford University provided expertise in whole artery immunohistochemistry and Itzel Llorente from Instituto de Fisiología Celular at UNAM performed cell culture and transfection of HEK293 cells with TRPV1. Thanks to Professor Christian Aalkjaer, University of Arhus for his insightful comments about sensory nerves and sodium hydrogen exchangers.

## DISCLOSURES

None

## AUTHOR CONTRIBUTIONS

EAF generated and analysed data, wrote the manuscript. MBA and KER generated electrophysiological data. KD supervised generation of whole artery staining. VB and ISC supervised the PCR, WB and ICC data and edited the manuscript. APA, TAJ, GWA and TR edited the manuscript. IAG supervised the projected, edited the manuscript and provided the funding for the project.

## Notes

### Competing Interest Statement

The authors have declared no competing interest.

